# Nucleotide excision repair is impaired by binding of transcription factors to DNA

**DOI:** 10.1101/028886

**Authors:** Radhakrishnan Sabarinathan, Loris Mularoni, Jordi Deu-Pons, Abel Gonzalez-Perez, Núria López-Bigas

## Abstract

Somatic mutations are the driving force of cancer genome evolution^1^. The rate of somatic mutations appears in great variability across the genome due to chromatin organization, DNA accessibility and replication timing^2-5^. However, other variables that may influence the mutation rate locally, such as DNA-binding proteins, are unknown. Here we demonstrate that the rate of somatic mutations in melanoma tumors is highly increased at active Transcription Factor binding sites (TFBS) and nucleosome embedded DNA, compared to their flanking regions. Using recently available excision-repair sequencing (XR-seq) data^6^, we show that the higher mutation rate at these sites is caused by a decrease of the levels of nucleotide excision repair (NER) activity. Therefore, our work demonstrates that DNA-bound proteins interfere with the NER machinery, which results in an increased rate of mutations at their binding sites. This finding has important implications in our understanding of mutational and DNA repair processes and in the identification of cancer driver mutations.

---

The accumulation of somatic mutations in cells results from the interplay of mutagenic processes, both internal and exogenous, and mechanisms of DNA repair. Recent efforts to sequence the whole genome of tumor samples from different tumor types^7,8^ have shed light on this interplay. On the one hand, mutational signatures associated to various tumorigenic mechanisms have been identified across cancer types^9^; on the other, genomic features such as chromatin organization, DNA accessibility, and DNA replication timing^2-5^ have been associated to the variation of somatic mutation rates at the megabase scale. Two recent studies proposed a causal relationship between the accessibility of chromosomal areas to the DNA repair machinery and their mutational burden. Supek and Lehner, 2015^10^ point to variable repair of DNA mismatches as the basis of the megabase scale variation of somatic mutation rates across the human genome. Polak et al. 2014^4^ attributed lower somatic mutation rates at DNase-I hypersensitive sites (DHS) in cell lines and primary tumors than at their flanking regions and the rest of the genome to higher accessibility to the global genome repair machinery. Similarly, nucleosome occupancy has been linked to regional mutation rate variation between the nucleosome bound DNA and linker regions^11-13^, while two recent studies found a relation between transcription factor binding sites (TFBS) and nucleotide substitution rates. Reijns et al. 2015^14^ detected increased levels of nucleotide substitutions around TFBS in the yeast genome, which was attributed to DNA-binding proteins acting as partial barriers to the polymerase-delta-mediated displacement of polymerase-alpha-synthesized DNA. Katainen et al. 2015^15^ found that CTCF/cohesin-binding sites are frequently mutated in colorectal tumors and in a small subset of tumors of other cancer types, and suggest that these mutations are probably caused by challenged DNA replication under aberrant conditions.

To elucidate the impact of DNA-binding proteins on DNA repair, we analyzed the somatic mutation rate at TFBS in the genomes of 38 primary melanoma samples sequenced by TCGA^16^. We found that the mutation rate was approximately five times higher in active TFBS, i.e., those overlapping DHS (Fig. 1a) than in their flanking regions (*P* < 2.2 × 10^−6^, chi-square test). We determined that this elevated mutation rate could not be explained by the sequence context (Fig. 1a), and that it did not occur at inactive TFBS (Fig. 1a and Extended Data Fig. 1), indicating that it is directly related to the protein bound to DNA. Furthermore, this enrichment for mutations appeared at the active binding sites of most transcription factors (TFs) (Fig. 1b, Extended Data Fig. 2 and Supplementary Table 1); the signal was discernible in most analyzed melanoma samples (Fig. 1c and Supplementary Table 2), and it increased with genome-wide mutation rate.

**Figure 1.**
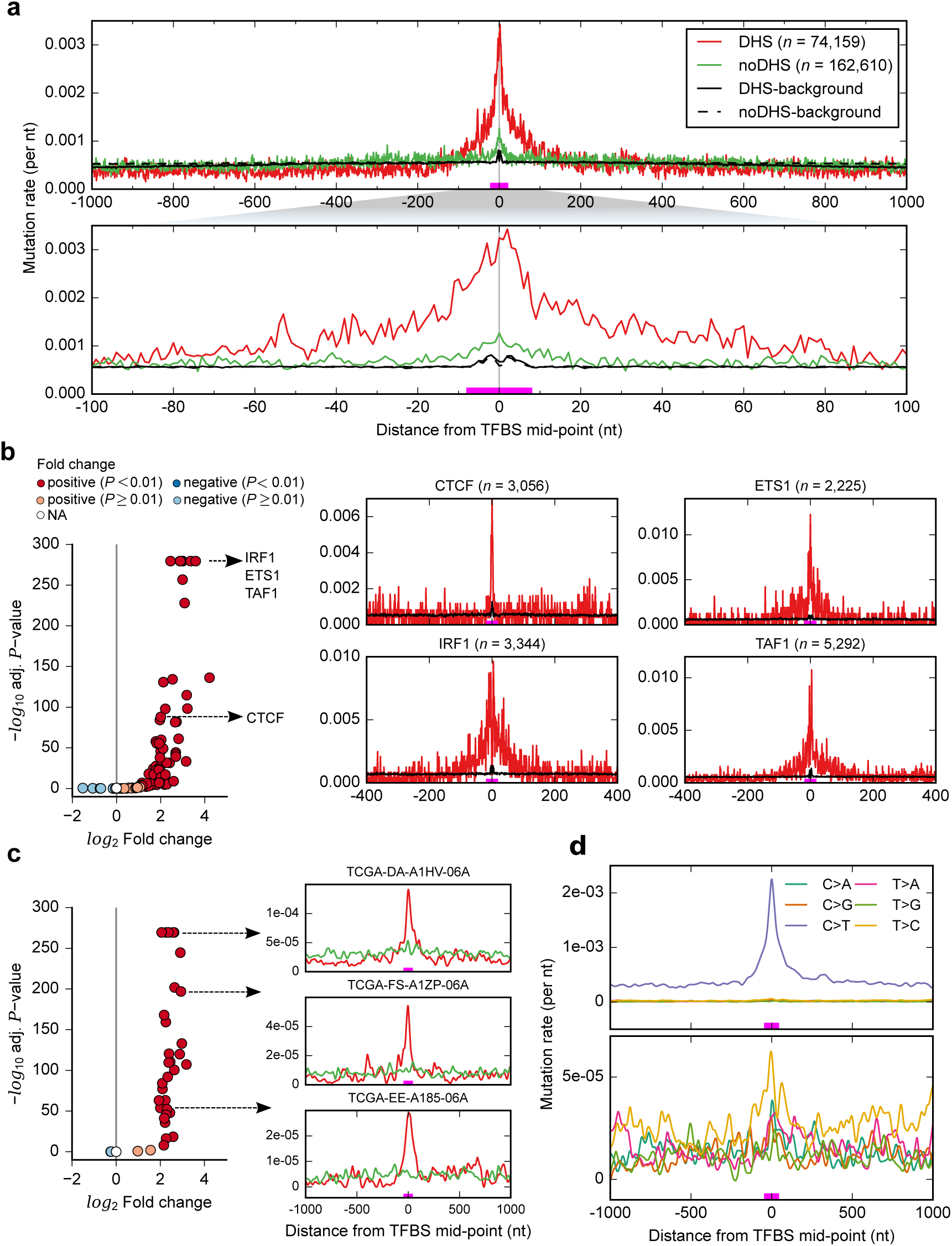
Elevated mutation rate at TFBS in melanomas. **a**, Mutation rates are approximately five-fold higher within active TFBS, those overlapping DHS in melanocytes, than in flanking regions (red line). In contrast, non-active TFBS, those non-overlapping DHS in melanocytes, do not show increased mutation rates (green line). The high increase in mutation rate is not explained by sequence context; black lines show the expected mutation rate per position when distributing all observed mu-tations in the region according to the probability of mutations in different trinucleotide contexts. **b**, A significant increase in mutation rate in TFBS compared to flanking regions is observed for most indi-vidual transcription factors and (c) in most of the individual melanoma samples. The log2 fold change (FC) on the *x*-axis represents if the mutation rate in TFBS is higher (positive FC) or lower (negative FC) than the expected, and the corresponding significance value (from chi-square test) is shown on the *y*-axis for each transcription factor. **d**, The contribution of C>T mutations to mutational density is higher compared to the other mutation types. The zero coordinate in the *x*-axis corresponds to the TFBS mid-point, and the magenta line above it represents the average size of TFBS.

**Figure 2.**
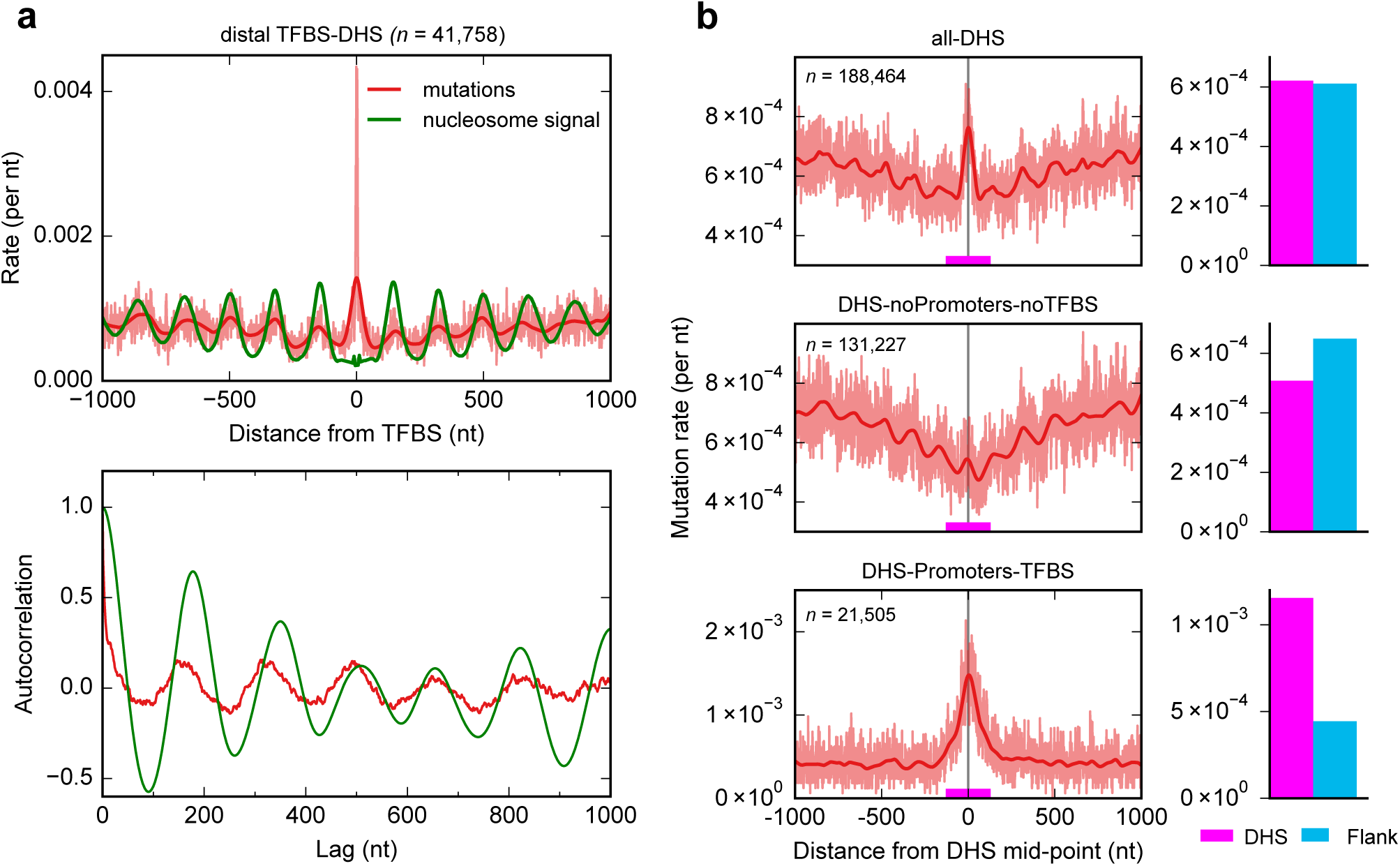
Mutation rate at distal TFBS and DHS sites. **a**, Mutation rate in distal TFBS, which are 5 kb away from transcription start sites. Similar to proximal TFBS as shown in Fig. 1a, the mutation rate is elevated at the center of core TFBS compared to the flanking. In addition, periodic peaks of mutation rate in the flanking regions of binding sites correlate well with the nucleosome positioning (green line). This is further supported by the Autocorrelation analysis (bottom panel) that shows the periodic peaks are observed at a distance of ~146bp, which coincides well with the size of the DNA being wrapped in nucleosomes, separated by short stretch of linker DNA. The zero coordinate in the *x*-axis corresponds to the TFBS mid-point. **b**, Mutation rate centered in DHS in melanomas is shown (top panel). In the subset of DHS regions outside promoter regions and non-overlapping TFBS (DHS-noPromoter-noTFBS), the peak in mutation rate disappears and only a valley is observed (middle panel). In the subset of DHS regions in promoters overlapping TFBS (DHS-Promoters-TFBS), the peak in mutation rate is more apparent. The actual mutation rate values are shown in light red and the best-fit spline is shown in dark red. The zero coordinate in the *x*-axis corresponds to the DHS peak mid-point, and the magenta line above it represents the average size of DHS (~150nts). The barplot at the right of each panel compares the mutation rate in the DHS and the flank for each group of regions.

Most somatic mutations in melanocytes are caused by exposure to ultraviolet (UV) radiation^9^. UV radiation causes specific DNA lesions or DNA photoproducts –cyclobutane pyrimidine dimers (CPDs) and (6-4) pyrimidine–pyrimidone photo-products ((6-4)PPs), at the sites of dipyrimidines^17^. As expected, C>T (G>A) mutations predominated over other nucleotide changes in melanomas (Fig. 1d), both within TFBS and at their flanks. This could be explained by either a faulty DNA repair or higher probability of UV induced lesions^18-19^ at protein-bound DNA.

Next, we focused on active TFBS in distal regions from transcription start sites, and again found increased mutation rate at binding sites, flanked by periodic peaks of mutation rate observed at a distance of ~146bp, which coincides well with the size of the DNA being wrapped in nucleosomes. When we superimposed the nucleosomes positioning signals from ENCODE^20^ and these mutation rate peaks, we verified that their positions matched perfectly (Fig. 2a). Furthermore, we found that the peak of mutation rate observed at the center of DHS regions occurred exclusively at TFBS located within promoter regions (DHS-Promoters-TFBS), and was absent from DHS-noPromoter-noTFBS (Fig. 2b). This corroborated that whatever the process causing the increment of mutation rate it required that the proteins be bound to the DNA.

We then inquired if the cause of the higher mutation rate in TFBS and nucleosomes was the reduced accessibility to the protein-bound DNA of the NER machinery. Non-repaired nucleotides would be by-passed by polymerases carrying out translesion DNA synthesis, thus resulting in mutations^21^. To test it we assembled nucleotide-resolution maps of the NER activity of the two products of UV-induced DNA damage, CPDs and (6-4)PPs, generated by Hu et al., 2015 using XR-seq in irradiated skin fibroblasts^6^. In XR-seq, the excised ∼30-mer around the site of damage generated during nucleotide excision repair is isolated and subjected to high-throughput sequencing. When we analyzed the genome-wide signal of this NER map, we found a strong decrease in the amount of CPD and (6-4)PP repair at the center of TFBS (Fig. 3a and Extended Data Fig. 3), compared to their flanking regions. The decrease was apparent both in wild-type cells (NHF1), and CS-B mutant cell lines, which lack transcription-coupled repair^6^ (Fig. 3a and Extended Data Fig. 3), and it appeared at the binding sites of individual transcription factors (Extended Data Fig. 4). Moreover, we found that the level of DNA excision repair (and the mutation rate) at TFBS correlated with the strength of their binding (Fig. 3B and Extended Data Fig. 5). We concluded from these observations that the higher mutation rate observed at active TFBS is caused by a decrease of the NER activity.

**Figure 3.**
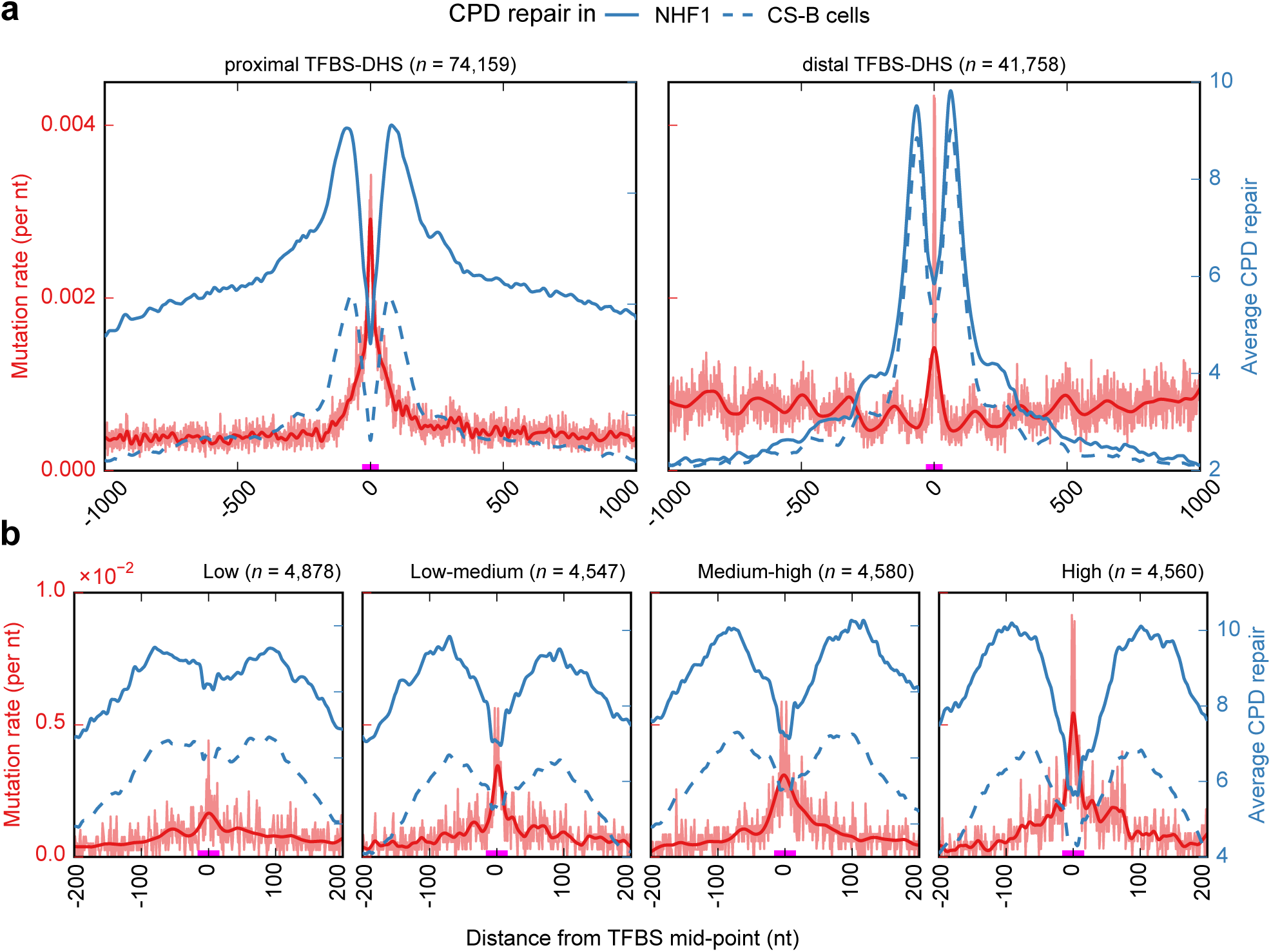
Regions around TFBS show a decrease in nucleotide excision repair. **a**, Mutation rate around TFBS is plotted (red line) alongside the average repair of UV-light induced DNA damage, cyclobutane pyrimidine dimers (CPD), in wild-type and CS-B mutant cell lines (blue line). A sharp decrease in nucleotide excision repair is evident at the core TFBS both in case of proximal and distal. **b**, The level of nucleotide excision repair (and the resulting mutation rate) in TFBS correlated with the strength of the binding of the transcription factor to its site. The binding sites were classified into four quartiles (Low to High) using the ChIP-seq read coverage that reflects the strength of binding or occupancy (as in Rejins et al., 2015^14^). The binding sites in the “High” quartile (last panel) show higher mutation rates at the center (correlating with the lower repair) compared to the “Low” quartile (first panel). The zero coordinate in the x-axis corresponds to the TFBS mid-point, and the magenta line above it represents the average size of TFBS.

**Figure 4.**
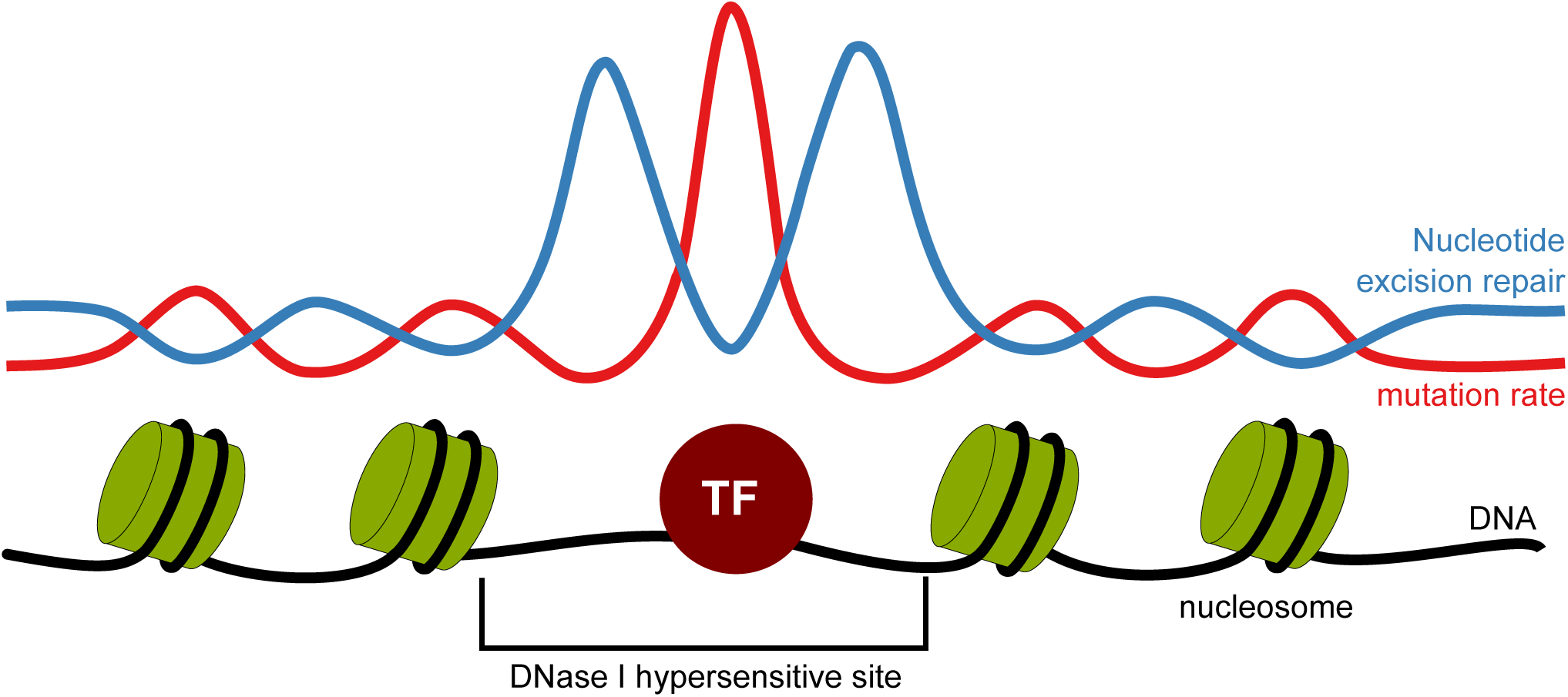
Model showing the mutation rate and repair rate in TFBS and nucleosome sites. The model shows that the accessibility of the DNA to the nucleotide excision repair (NER) machinery directly determines the distribution of mutational density at the nucleotide scale. Lower NER activity is observed at the TFBS bound region (within DHS region) and the nucleosome positions in the flank, compared to the nucleosome free regions (DHS and linkers). This NER activity corroborates the observed high mutation rate in transcription factor and nucleosome bound regions.

A previous study related higher DNA repair activity at DHS than that outside DHS to greater accessibility to the repair machinery^4^. By specifically deconvoluting the signal of mutation rate within DHS, our work goes a step beyond to show that bound TFs at the center of DHS actually hinder DNA repair. This interplay of greater NER at DHS and lower NER at TF bound sites in their center results in a volcano-shaped pattern of NER activity around the TFBS, with a strong depletion exactly at its center flanked by two mountains in the DHS area around it (Fig. 3). The volcano shape is more pronounced at distal TFBS, those that occur distant from transcription start sites (TSS) (Fig. 3a), which may be explained by the presence of shorter regions of open chromatin surrounded by compacted DNA. Moreover, a periodicity in NER activity is observable for the first nucleosomes around TFBS (Fig 3a), which matches nicely the previously noted periodical variation of the mutation rate. Also in coherence with the mutation rate pattern, the signal of decreased NER activity is clearer at the center of DHS-Promoters-TFBS, exactly at the position of the TFBS (Extended Data Fig. 6c). These results demonstrate that repair activity in DHS regions is in general higher than in non-DHS regions, supporting previous observations^4^, however this activity is specifically impaired at sites with bound transcription factors.

NER consists of two pathways: global repair –targeting the lesions in a genome-wide manner– and transcription-coupled repair that recognize lesion within transcribed regions^17^. These pathways differ in the initial steps of damage recognition, although they share the core component that excise damaged regions. To discern the effect of DNA bound TFs on transcription coupled NER we focused on transcribed regions centered at TFBS at least 200 bps downstream of TSS, and plotted together mutation rate and XR-seq data in XP-C cells, which only have transcription-coupled repair^6^. Mutation rate is also increased at the center of transcribed TFBS, and the volcano shape of repair rate in XP-C cells is apparent for TFs bound to either template or non-template strand (Extended Data Fig. 7). This result demonstrates that the decrease in NER caused by bound TFs results from impairment of both NER pathways.

NER recognizes and repairs other DNA lesions beside those induced by UV light, such as DNA adducts induced by smoking-related carcinogens (e.g. benzo[*a*]pyrene diol epoxide)^22^. We therefore hypothesized that the conclusion we had drawn from the observations made in melanomas could be extended to other tumor types. We observed higher mutation rates at TFBS in lung adenocarcinomas and lung squamos cell carcinomas, in particular for C>A variants, which correspond to the mutations caused by tobacco smoking^9^ (Extended Data Fig. 8). In contrast, no increment of the mutation rate in TFBS is observed in colon adenocarcinomas, where NER activity is not expected to play a major role in the mutational process.

Two previous studies have described abnormal mutation rates in connection with a group of DNA bound TFs in yeast^14^ and CTCF/cohesin sites in a subset of colorectal tumors^15^. However, in contrast to our results, in neither of these studies the peaks of mutation rate were caused by impairment of NER resulting from bound proteins. In the former, higher mutation rate at specific TFBS were related to polymerase-delta-mediated displacement of polymerase-alpha-synthesized DNA during replication. In the latter, higher mutations at CTCF/cohesin sites of a subset of colorectal tumors, was attributed to challenged DNA replication under aberrant conditions.

In summary, our results demonstrate that the accessibility of the DNA to the NER machinery directly determines the distribution of mutational density at the nucleotide scale. The increased repair in freely accessible, nucleosome-free, DNA around TFBS and the decline in repair efficiency exactly at TFBS produces a lower mutation rate in the periphery of DHS sites and higher mutation rate at their center (Fig 4). Moreover, periodic signals of higher mutation rate and lower NER in close chromatin regions coincide with nucleosome occupancy, suggesting that nucleosomes produce the same type of impairment to NER.

These findings have strong implications for our basic understanding of the mechanisms of DNA repair in human cells, as well as for the study of tumor evolution and cancer-associated somatic mutations. They indicate that most mutations in TFBS accumulate due to faulty repair at these sites. Therefore, methods designed to identify potential somatic driver mutations, in non-coding regions, which typically exploit the mutational patterns of genomic elements must construct models of the background mutation rate that accurately take into account the increased mutation density at TFBS due to faulty repair.

## Methods

### Mutation data

Whole-genome somatic mutations of 38 skin cutaneous melanomas (SKCM), 46 lung adenocarcinomas (LUAD), 45 lung squamous cell carcinomas (LUSC), and 42 colorectal adenocarcinomas (CRC) identified by TCGA were obtained from Fredriksson et al., 2014^16^. As suggested by the authors of that paper, we considered in our analyses only single nucleotide substitutions with a minimum variant frequency of 0.2 and which do not overlap dbSNP entries (v138). The total number of mutations of each cancer type passing these thresholds is listed in Extended Data Table 1. We separated CRC samples into two groups: hypermutated (with mutations of the DNA polymerase epsilon (POL-E) gene; *n* = 8 samples) and hypomutated (the rest; n = 34 samples).

### Genomic elements

The genomic coordinates of transcription factor binding sites (TFBS), i.e., TF motif match under ChIP-seq peak regions, were obtained from ENCODE^20^. These comprised the binding sites of 109 transcription factors (TF) as used in Khurana et al., 2013^23^. We also obtained from ENCODE predicted binding sites of 52 transcription factors which are not supported by ChIP-seq peaks (termed unbound TFBS). In addition, we obtained the binding sites of 32 TFs used in Reijns et al., 2015^13^. We treated the latter as an independent data set, and following the authors of the original paper,^13^ we clustered the TFBS into quartiles according to the binding strength or occupancy of the TFs to their sites – quantified through ChIP-seq read coverage.

As promoters, we considered the DNA sequences up to 2.5kb upstream of transcription start sites (TSS) of all protein coding genes in GENCODE^24^ (v19). Promoter regions overlapping coding sequences (CDS) or untranslated regions (UTRs) were excluded. We classified TFBS as either proximal –i.e., overlapping these upstream promoters– or distal –i.e., those located in intergenic regions, with no annotated TSS (as per GENCODE v19) within 5kb distance on either side. A third group of TFBS was composed of those located downstream TSS (between +200bp and +500bp) and which do not overlap with the upstream 2.5kb promoter regions –i.e., TFBS in transcribed regions.

All TFBS overlapping DNase I Hypersensitive sites (DHS) identified by the Epigenome roadmap project^25^ in primary cell types most closely matching the cell of origin of each tumor type (see below) were considered active. We considered only DHS sites identified by the Hotspot algorithm (narrowPeaks in FDR 1%), which are typically 150nts long. For each cancer type, the matching primary cell type was selected based on the recent study by Polak et al., 2015^5^ (Extended Data Table 1). We chose the DHS from primary cell types (from Epigenome Roadmap project) instead of cell lines (from ENCODE), because the chromatin features of the cell of origin of a tumor has been shown to correlate better with its mutation profile than that of matched cancer cell lines^5^. However, we selected the TFBS detected by ENCODE in cell lines (see above) due to the lack of TF binding site annotations in primary cells analyzed by the Epigenome Roadmap project^25^.

We then classified the TFBS in the samples of each tumor type as active or inactive based on their overlap, or lack thereof, with DHS regions (minimum 1bp) of the matched primary cell type. Unbound TFBS (see above), which do not overlap with TF peaks or DHS regions, were considered as inactive TFBS and used as negative control to compare with the active TFBS (in Extended data Fig. 1). All genomic co-ordinates of TFBS used in this study as part of any aforementioned category are available at http://bg.upf.edu/tfbs.

### Mutation rate estimation

In order to compare the mutation rate in TFBS to their neighboring regions, we considered flanking stretches of 1000 nucleotides at both sides of the TFBS mid-point. To exclude regions that could bias the mutation rate analyses, prior to mapping the somatic mutations to these selected 2001 nts windows, we filtered out: any regions overlapping a) coding sequences, and b) UCSC Browser blacklisted regions, often misaligned to sites in the reference assembly, (Duke and DAC) and low unique mappability of sequencing reads (“CRG Alignability 36' Track”^26^, score < 1) (http://genome.ucsc.edu/cgi-bin/hgFileUi?db=hg19&g=wgEncodeMapability). In addition, regions that overlap other TFBS within flanking regions (immediately upstream or downstream the TFBS) were excluded. The resulting filtered windows of each TFBS were then aligned (taking as reference the TFBS centers), and the mutation rate of every column *i* within the window was calculated as the total number of mutations mapped to nucleotides in column *i* divided by the total number of nucleotides observed in column *i* (after filtering). We computed this mutation rate for each TF separately, as well as globally for all TFs. In the latter case, prior to the calculation, we removed any repeated chromosomal positions (from different TFs) observed in a column.

In the case of the analysis center on DHS, we considered flanking stretches of 1000 nucleotides at both sides from DHS peak center and followed the same steps mentioned above to filter mutations and to compute the mutation rate.

### Background mutation rate estimation

In order to check if the mutation rate observed at each position was expected due to the local sequence context, we randomly introduced the same number of mutations observed at each window following the probability of occurrence of each mutation according to its tri-nucleotide context. We computed the probability of occurrence of all possible 96 tri-nucleotide changes in each cancer type based on the total number of observed mutations in all its samples. We also computed separate probabilities of occurrence of all 96 tri-nucleotide in active and inactive TFBS from the mutations observed in each category. The mutation rate of each randomly generated set of changes, was computed for each column as explained above. This procedure was repeated 1000 times to compute the mean random mutation rate of every column in the motif.

### Enrichment analysis

To identify if TFBS is enriched for mutations compared to the immediate flanking region, we compared the ratio of the total number of mutations to the total number of nucleotide positions within the TFBS region (-15 to 15nts) and that of the flanking region (16 to 1000nts on either side) using a chi-squared test. We performed this test for all transcription factors and for each individual tumor, and corrected the resulting*p*-values for multiple-testing using the Benjamini-Hochberg procedure^28^. In addition, we computed the fold change of mutation rates through the expected frequencies obtained from chi-squared tests. Both, the fold change and adjusted *p*-values are shown in Figure 1b-c.

### Nucleotide excision repair data

The genome-wide maps of nucleotide excision repair of two types of UV-induced damage, cyclobutane pyrimidine dimers (CPD) and (6-4) pyrimidine-pyrimidone photoproducts ((6-4)PP), available for three different cell lines –i) wild-type NHF1 skin fibroblasts, ii) XP-C mutants, lacking the global repair mechanism, and iii) CS-B mutants lacking transcription-coupled repair– were obtained from Hu et al., 2015^6^. The dataset contains normalized read counts for fixed steps of 25bp across the genome, for the forward and reverse strands separately. We kept these for our analyses and also generated strand independent data as the average of normalized read counts from both strands for every nucleotide position. These average read counts were mapped to the TFBS centered windows (2001bp), filtered and aligned to the TFBS mid-point as described above. We computed the average repair rate for each column *i* of these windows as the total number of average read counts mapped to the nucleotides in the column *i* divided by the total number of nucleotides in the column i, as described above for the mutation rate.

### Nucleosome signals

Genome-wide nucleosome positioning signals (density graph) of ENCODE cell line GM12878 (lymphoblastoid cell line) were downloaded *via* the UCSC genome browser (http://hgdownload.cse.ucsc.edu/goldenpath/hg19/encodeDCC/wgEncodeSydhNsome/). We then mapped them to the TFBS centered windows, and similar to mutation and repair rates, we computed the average signal per column *i* of the window as the sum of signal values mapped to the nucleotides in column *i* divided by the total number of nucleotides in column *i.*

### Computational and statistical tools

BEDTools utilities^29^ were used to carry out operations as extensions or overlaps in the various analyses of genomic features (TFBS/DHS), as well as to map somatic mutations to genomic features. All curve fittings shown in figures (best-fit spline) were performed using the smooth.spline function from R^30^ (v3.0). The auto-correlation was performed using the *acf* function from statsmodels python package (http://statsmodels.sourceforge.net/).

## ACKNOWLEDGEMENTS

We acknowledge funding from the Spanish Ministry of Economy and Competitiveness (grant number SAF2012–36199), the Marató de TV3 Foundation, and the Spanish National Institute of Bioinformatics (INB). R.S. is supported by an EMBO Long-Term Fellowship (ALTF 568–2014) co-funded by the European Commission (EMBOCOFUND2012, GA-2012–600394) support from Marie Curie Actions. A.G.-P. is supported by a Ramón y Cajal contract.

## Supplementary information

**Supplementary Table 1:** Results of mutation rate enrichment at the binding sites of individual TFs in melanoma.

**Supplementary Table 2:** Results of sample-wise analysis of mutation rate enrichment at the active TFBS for 38 melanoma samples.

**Extended Data Figure 1:**
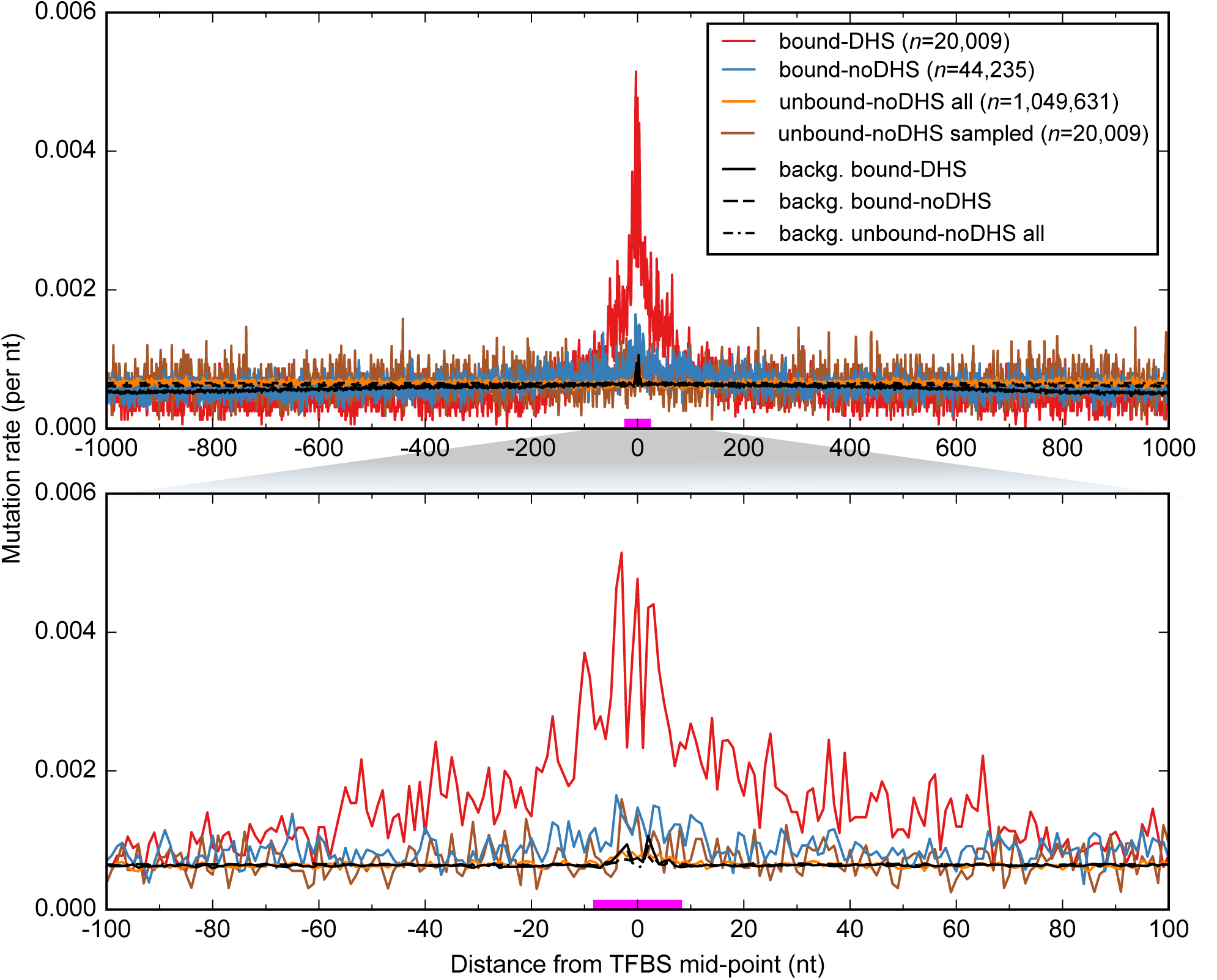
Higher mutation rate at bound TFBS compared to unbound TFBS in melanoma. The mutation rate is higher in active TFBS (bound by their TF and overlapping DHS; bound-DHS, red line) compared to: i) inactive TFBS (not overlapping any DHS; bound-noDHS, blue line); and ii) unbound inactive TFBS (not bound by TF and not overlapping any DHS; unbound-noDHS, orange line). The binding sites considered here correspond to the subset of TFs (n = 58) for which both the bound and unbound motif predictions are available from the ENCODE integrative analysis^20^. For comparison purposes, we sampled an equal number of unbound-noDHS TFBS (unbound-noDHS samples, brown line) as in the set of bound-DHS, and confirmed that the mutation rate is still higher in the bound TFBS. The background mutation rates of each group are represented as black lines. The zero coordinate in the *x*-axis corresponds to the TFBS mid-point, and the magenta line above it represents the average size of TFBS.

**Extended Data Figure 2:**
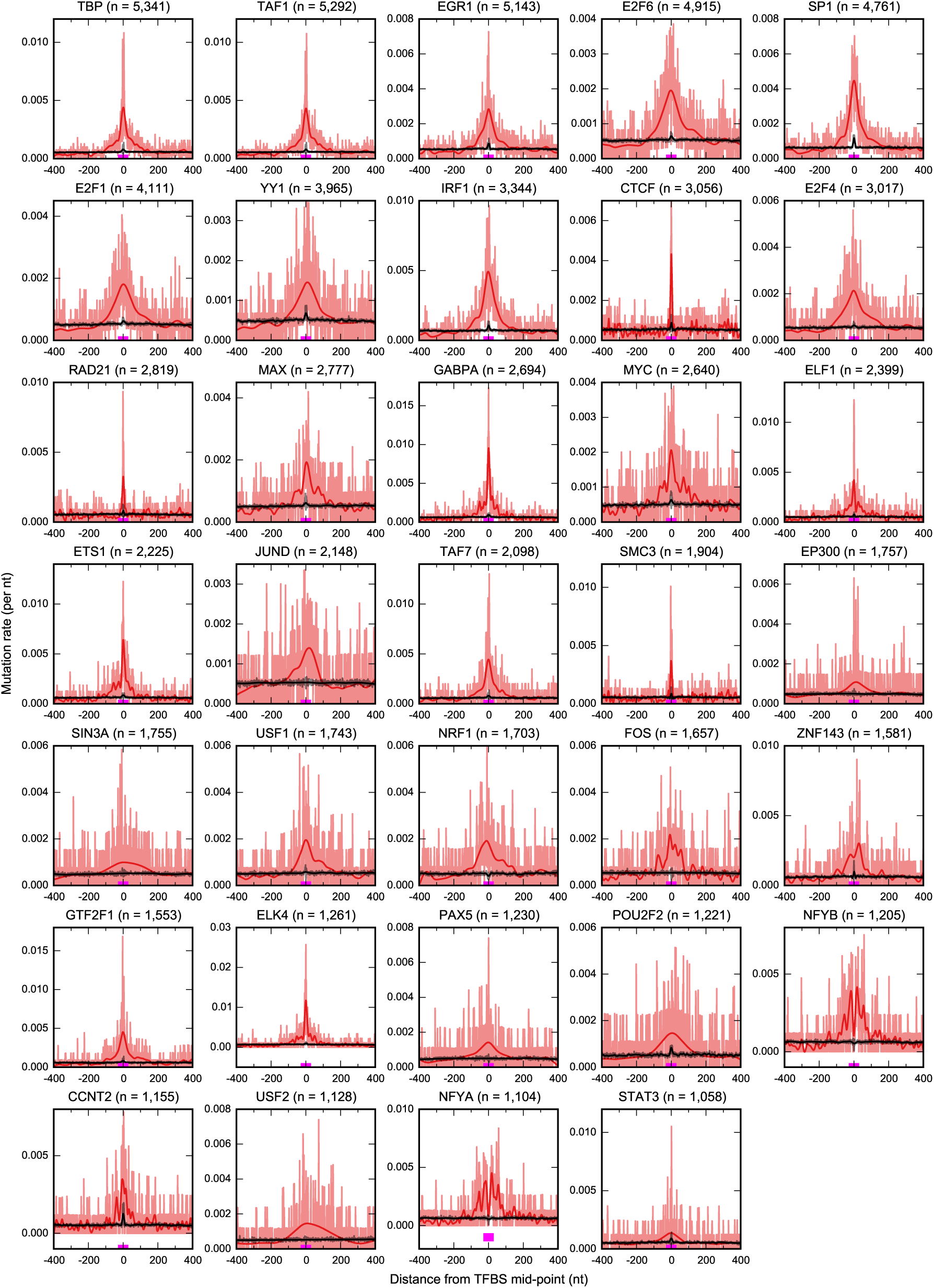
Elevated mutation rate at the binding sites of individual transcription factors (TF) in melanoma. Here, we show the mutation rate of the TFBS of all TFs with at least 1000 binding sites overlapping melanocytes DHS. The observed mutation rate is shown in red (light color in the background corresponds to the actual data points, and the thick solid line on top is the best-fit spline), while the background mutation rate is represented by the black line. The zero coordinate in the *x*-axis corresponds to the TFBS mid-point, and the magenta line above it represents the average size of TFBS.

**Extended Data Figure 3:**
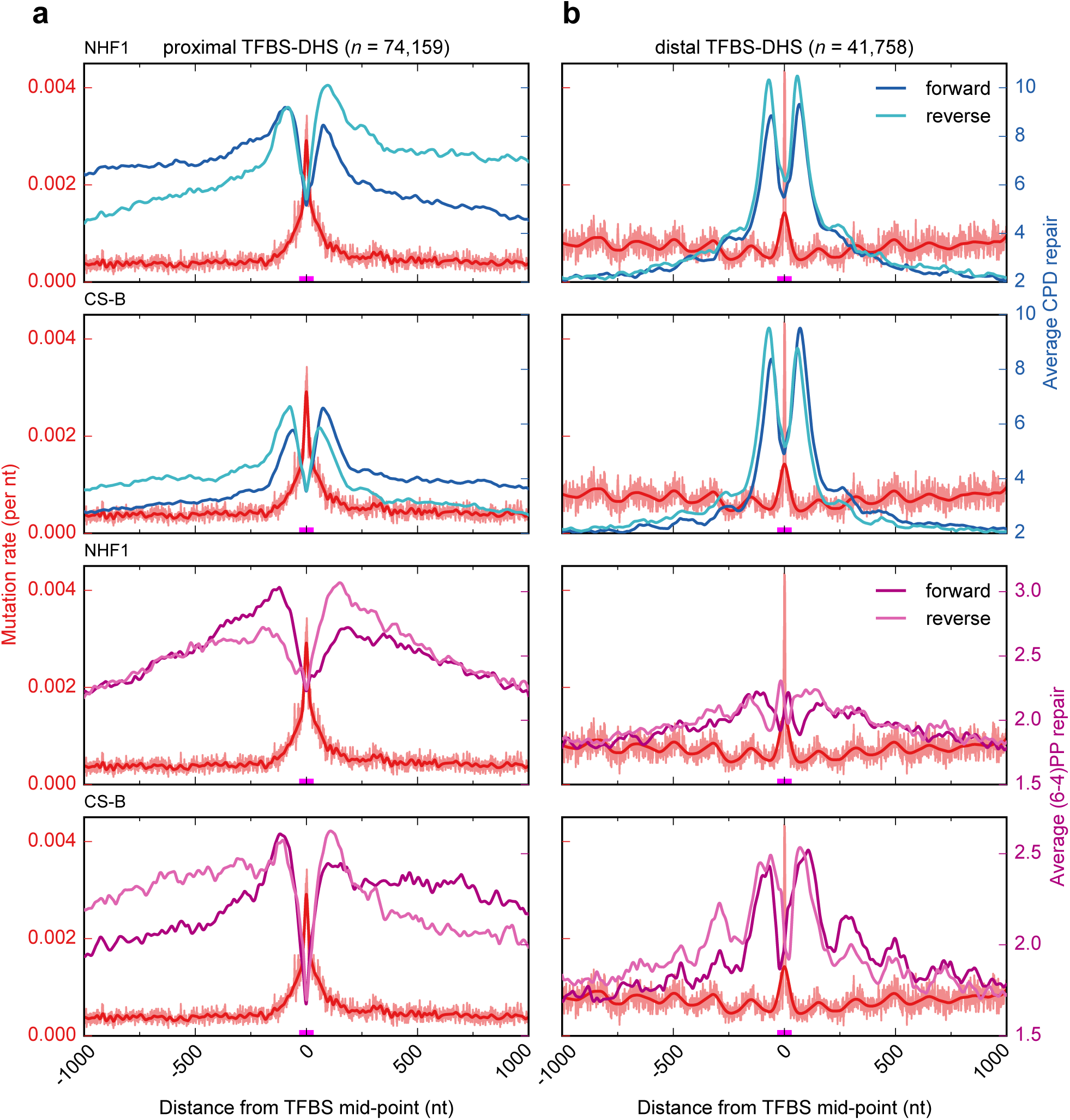
Regions around TFBS show a decrease in nucleotide excision repair. Mutation rate around TFBS plotted alongside the average repair of two types of UV-light induced DNA damage-cyclobutane pyrimidine dimers (CPD) and (6–4) pyrimidine-pyrimidone photo-products ((6–4)PP) in wild-type NHF1 cell line of skin fibroblasts and the CS-B mutant cell line for proximal (a) and distal (b) TFBS. The two top rows show the CPD repair on NHF1 and CS-B cells, respectively and the two bottom rows show the (6–4)PP repair in NHF1 and CS-B cells, respectively. Here, the average repair levels are shown separately for the forward and reverse strands of the genome (as provided by Hu et al., 2015^6^). The left *y*-axis corresponds to the mutation rate, and the right *y*-axis corresponds to the average repair rate. The zero coordinate in the *x*-axis corresponds to the TFBS mid-point, and the magenta line above it represents the average size of TFBS.

**Extended Data Figure 4:**
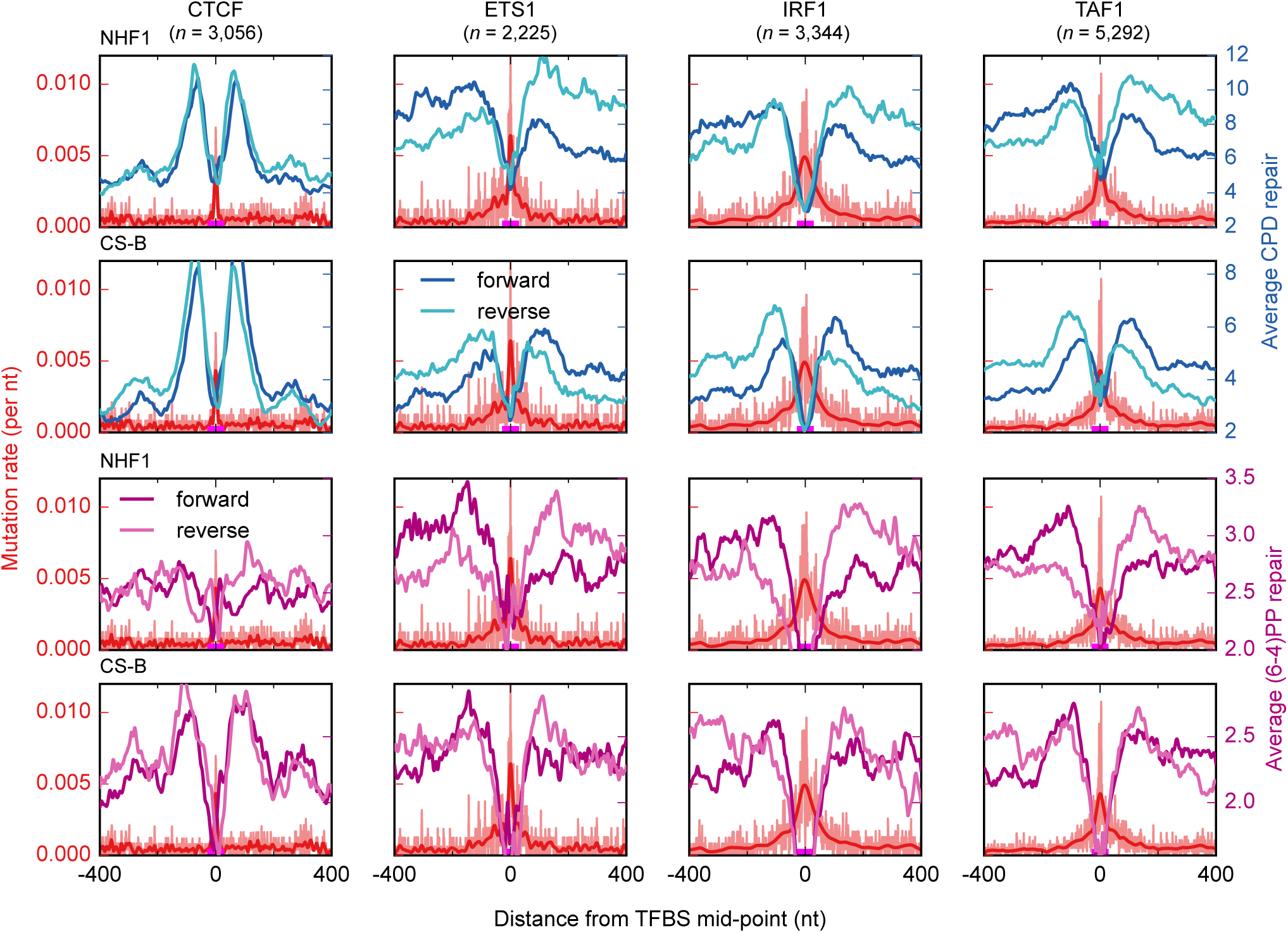
Lower level of nucleotide excision repair at binding sites of individual transcription factors. A lower level of nucleotide excision repair is observed at the binding sites of individual transcription factors. In the figure we present the numbers of CTCF, ETS1, IRF1 and TAF1in columns. The observed mutation rate is shown in red (light color in the background corresponds to the actual data points, and the thick solid line on top is the best-fit spline). The two top rows show the CPD repair on NHF1 and CS-B cells, respectively and the two bottom rows show the (6–4)PP repair on NHF1 and CS-B cells, respectively. Here, the average repair levels are shown separately for the forward and reverse strands of the genome (as provided by Hu et al., 2015^6^).

**Extended Data Figure 5:**
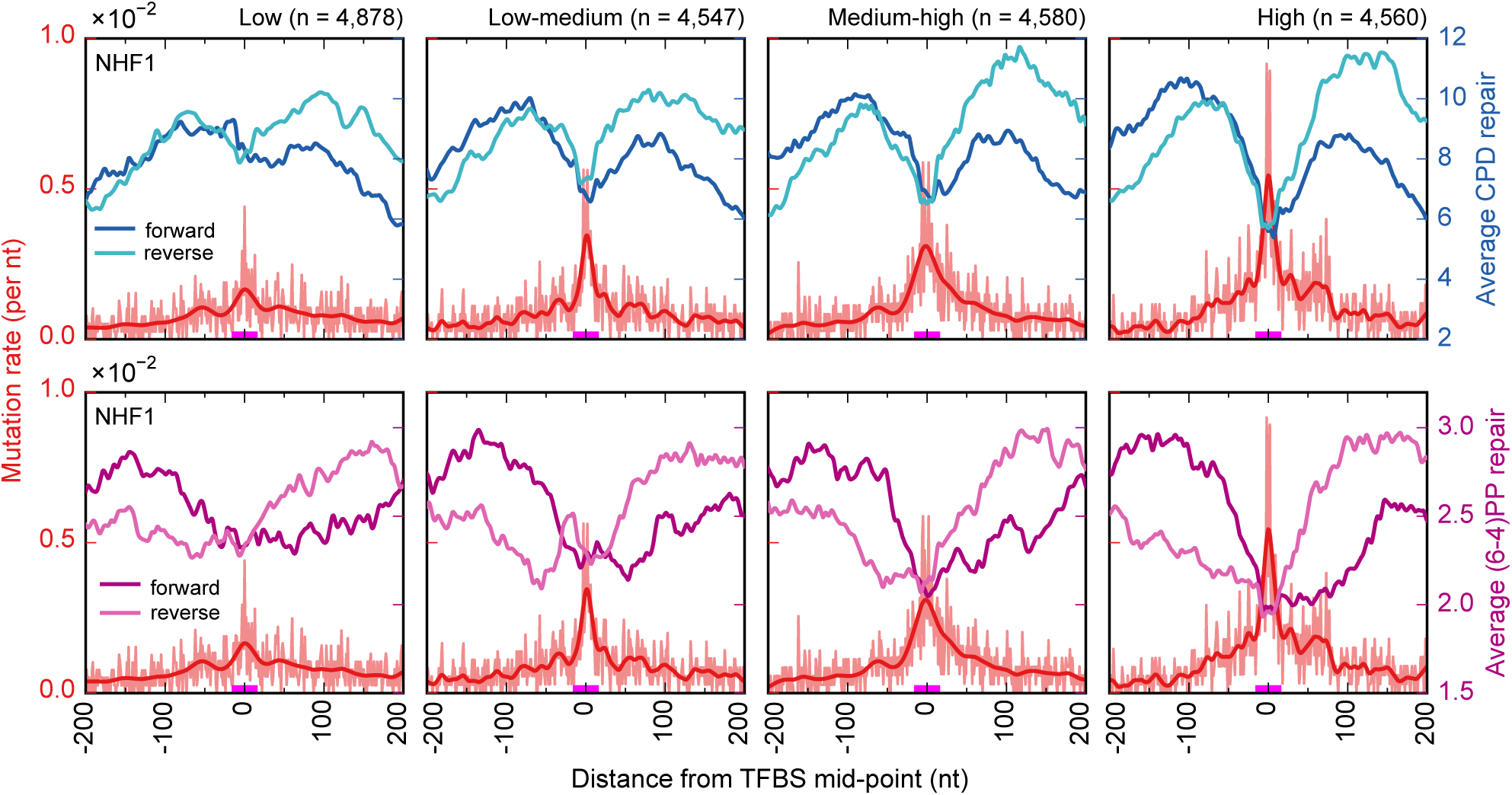
The level of nucleotide excision repair, and the resulting mutation rate in TFBS correlate with the strength of the binding signal of transcription factors to their sites. Regions around TFBS sites were obtained from Rejins et al., 2015^14^. As in ref. 14, the binding sites were classified into four quartiles (Low to High) using the ChIP-seq read coverage that reflects the strength of binding or occupancy. The binding sites in the “High” quartile (fourth column) tend to bear higher mutation rates at the center (correlating with lower repair) compared to the “Low” quartile (first column). The nucleotide excision repairs of two photoproducts (CPD and (6–4)PP) shown here are from NHF1 wild-type cell line. Average repair levels are shown separately for the forward and reverse strands of the genome (as provided by Hu et al., 2015^6^).

**Extended Data Figure 6:**
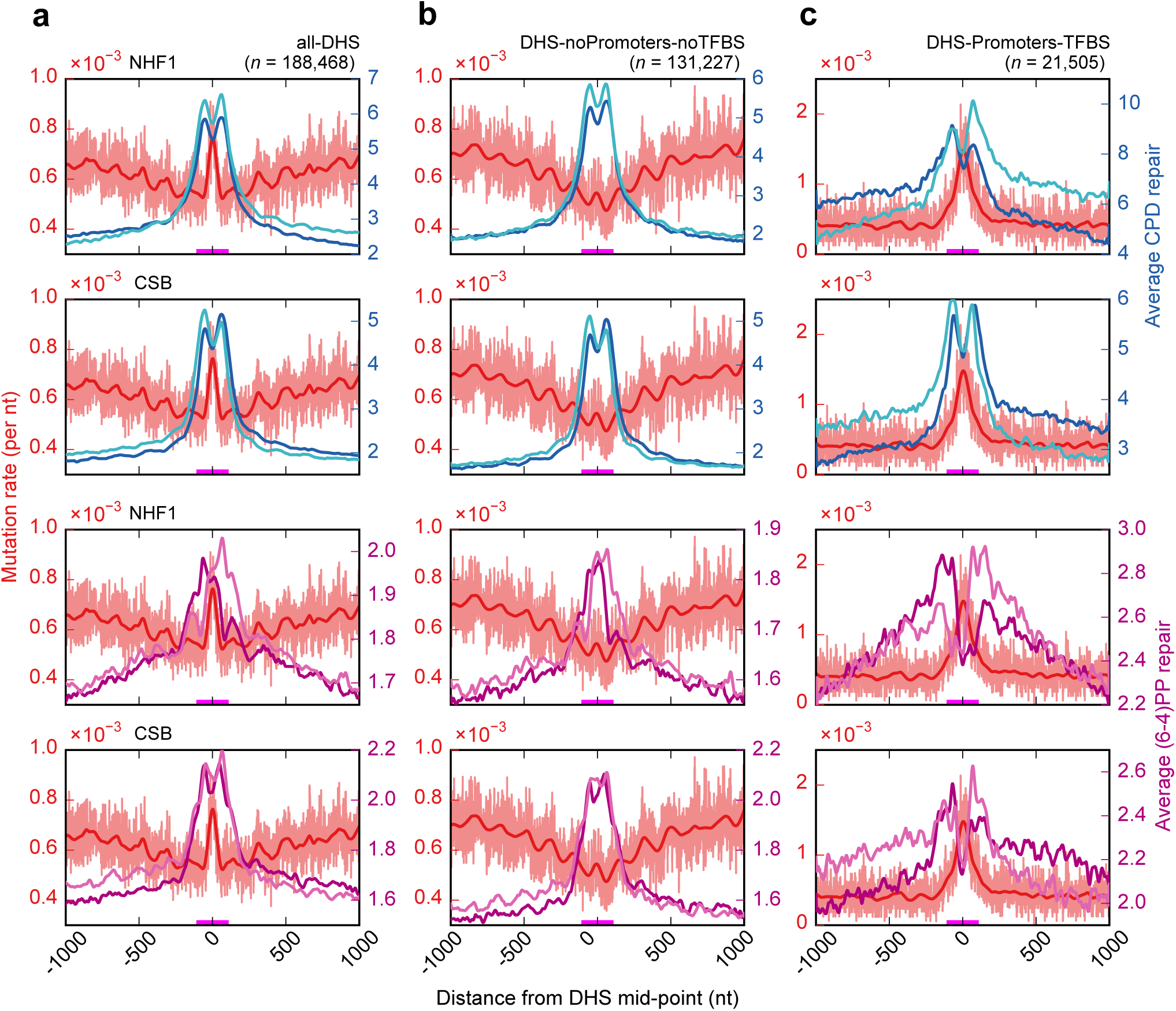
Nucleotide excision repair and mutation rate in DHS centered regions. The distribution of nucleotide excision repair in all DHS, DHS-noPromoter-noTFBS and DHS-Promoters-TFBS regions for the two types of UV-light induced DNA damages. The two top rows show the CPD repair on NHF1 and CS-B cells, respectively and the two bottom rows show the (6–4)PP repair on NHF1 and CS-B cells, respectively. Here, average repair levels are shown separately for the forward and reverse strands of the genome (as provided by Hu et al., 2015^6^). The zero coordinate in the x-axis corresponds to the DHS peak mid-point, and the magenta line above it represents the average size of DHS (~150nts).

**Extended Data Figure 7:**
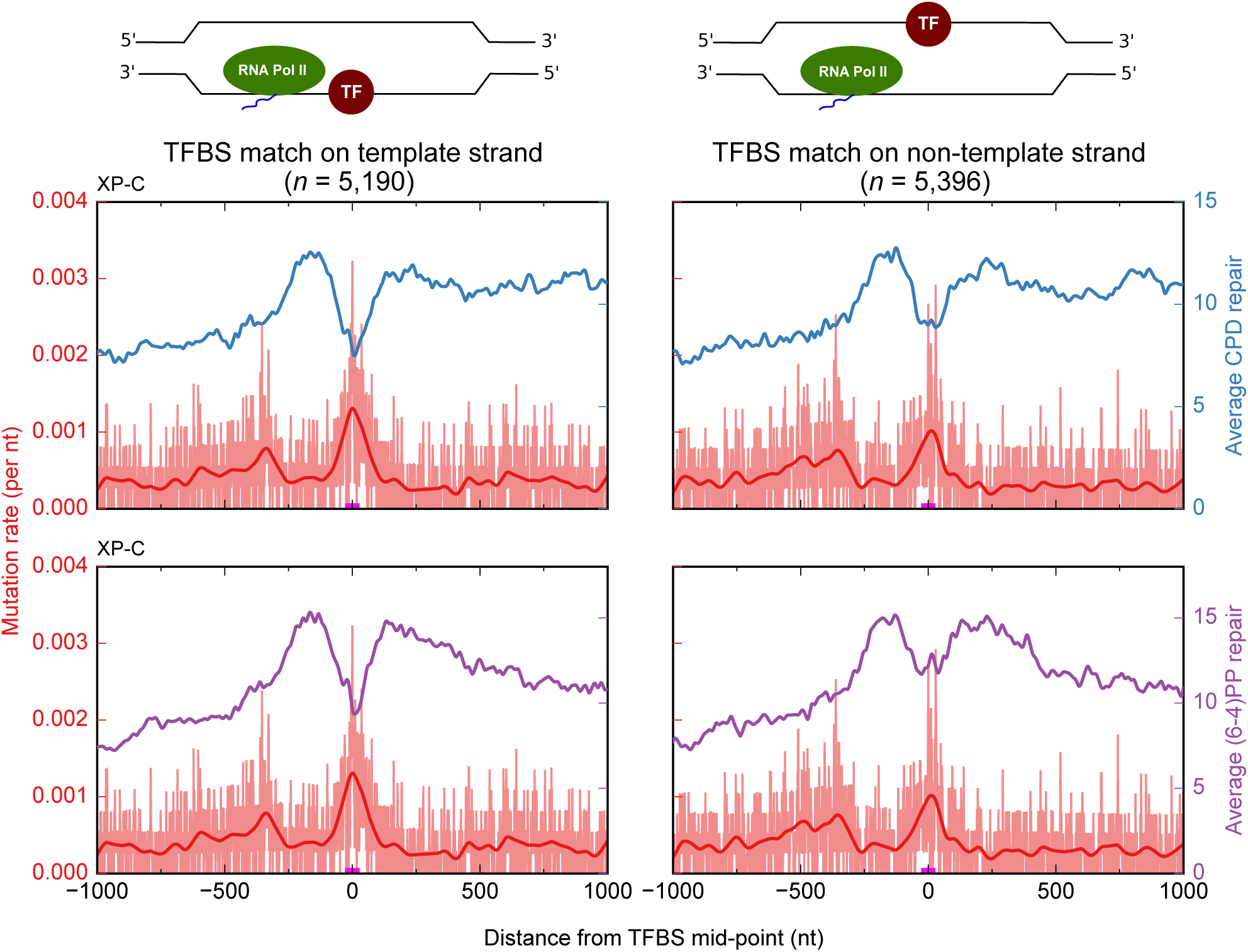
Transcription coupled-repair is impaired at active TFBS. To carry out this analysis, TFBS overlapping transcribed regions (located 200–500bp downstream of TSS) were centered at the TFBS mid-point. We plot the mutation and repair rates of UV induced damages (CPD and (6–4 PP)) in XP-C cells, which possess only transcription coupled repair capability. TFBS in either strand were separated: those in the template strand of the gene are shown in the left panel, while those in the non-template strand are presented in the right panel. All TFBS and their flanking regions are shown in the same orientation (5’ to 3’). This result shows that TF binding to both strands results in lower transcription-coupled NER activity.

**Extended Data Figure 8:**
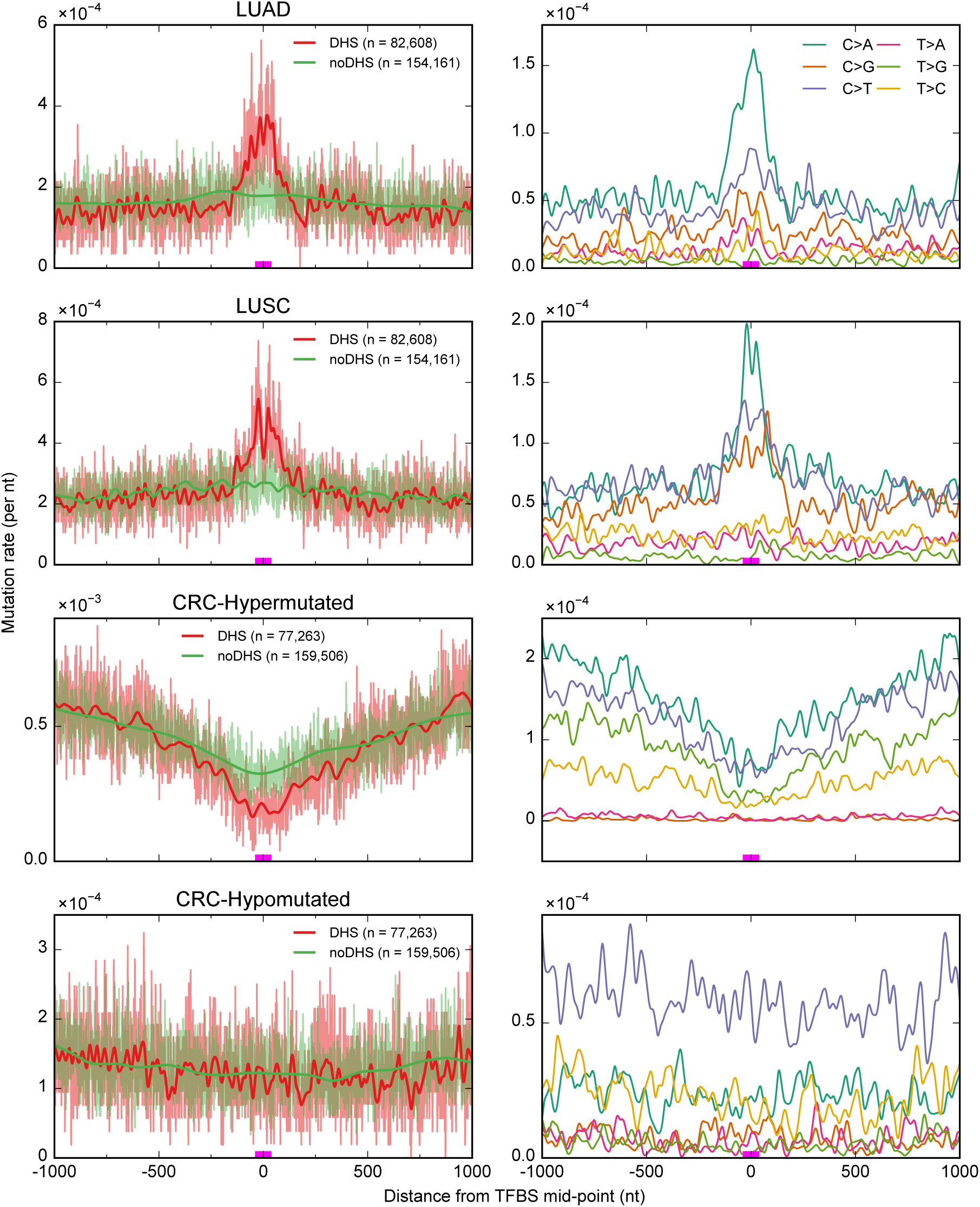
Mutation rate around TFBS in other cancer types. Mutation rates around TFBS of promoter regions of Lung Adenocarcinoma (LUAD), Lung Squamous Cell Carcinoma (LUSC), and Colorectal Cancer (CRC) are shown. CRC samples are separated into two groups, those with missense mutations of the DNA polymerase epsilon (POL-E) gene or Hypermutated (n = 8 samples) and the rest or Hypomutated (n = 34 samples). In the left column, the mutation rate is shown for active TFBS that overlap DHS sites (red line) and inactive TFBS that do not overlap DHS (green line). The right column graphs present the mutation rate of six different changes separately in active TFBS. In lung cancers (LUAD and LUSC), C>A changes, caused by tobacco carcinogens, contributes more to the elevated mutation rate, which indicates that NER activity is lower at these active TFBS.

**Extended Data Table 1.** The whole genome sequencing data of different cancer types from TCGA and the matching primary cell types from Epigenome roadmap.

| Cancer Type | Number of whole genome samples | Total number of mutations<sup>a</sup> | Mutation per tumor Median<sup>b</sup> | Matching primary cell type from Epigenome roadmap (EID)<sup>c</sup> |
| --- | --- | --- | --- | --- |
| Skin Cutaneous Melanoma (SKCM) | 38 | 3,336,384 | 46,600 | Foreskin Melanocyte Primary Cells (E059) |
| Lung adenocarcinoma (LUAD) | 46 | 1,030,242 | 14,166 | Fetal lung (E088) |
| Lung Squamous cell carcinoma (LUSC) | 45 | 1,404,152 | 32,357 | Fetal lung (E088) |
| Colorectal (CRC) | 42 | 3,556,383 | 11,594 | Fetal Intestine Large (E084) |
| CRC - *Hypermutated* | 8/42 | 2,909,900 | 404,986 | Fetal Intestine Large (E084) |
| CRC – *Hypomutated* | 34/42 | 646,483 | 9,792 | Fetal Intestine Large (E084) |

<sup>a,b</sup> The somatic mutations were previously called by Fredriksson et al., 2014, and the numbers presented here correspond to the total number of mutations after filtering as suggested by authors (see Materials section for more details). In the case of colorectal cancer (CRC), the samples were divided into two groups:
*Hypermutated and Hypomutated*, based on the presence of missense mutations in the DNA polymerase epsilon gene or not.
<sup>c</sup> For each primary cell type, the respective DHS identified by Hotspot algorithm (FDR 1%) were download from http://egg2.wustl.edu/roadmap/data/byFileType/peaks/consolidated/narrowPeak/

